# CLigopt: Controllable Ligand Design Through Target-Specific Optimisation

**DOI:** 10.1101/2024.03.15.585255

**Authors:** Yutong Li, Pedro Henrique da Costa Avelar, Xinyue Chen, Li Zhang, Min Wu, Sophia Tsoka

## Abstract

**Motivation:** Key challenge in deep generative models for molecular design is to navigate random sampling of the vast molecular space, and produce promising molecules that compromise property controls across multiple chemical criteria. Fragment-based drug design (FBDD), using fragments as starting points, is an effective way to constrain chemical space and improve generation of biologically active molecules. Furthermore, optimisation approaches are often implemented with generative models to search through chemical space, and identify promising samples which satisfy specific properties. Controllable FBDD has promising potential in efficient target-specific ligand design.

**Results:** We propose a controllable FBDD model, CLigOpt, which can generate molecules with desired properties from a given fragment pair. CLigOpt is a Variational AutoEncoder-based model which utilises co-embeddings of node and edge features to fully mine information from molecular graphs, as well as a multi-objective Controllable Generation Module to generate molecules under property controls. CLigOpt achieves consistently strong performance in generating structurally and chemically valid molecules, as evaluated across six metrics. Applicability is illustrated through ligand candidates for hDHFR and it is shown that the proportion of feasible active molecules from the generated set is increased by 10%. Molecular docking and synthesisability prediction tasks are conducted to prioritise generated molecules to derive potential lead compounds.

**Availability and Implementation:** The source code is available via https://github.com/yutongLi1997/CLigOpt-Controllable-Ligand-Design-through-Target-Specific-Optimisation.

## 1 Introduction

The development of novel drugs is dependent on the efficient discovery of novel molecules [1]. A critical challenge in this process is the difficulty in navigating the immense molecular space. Traditional methods, such as high-throughput screening, are inefficient as they require large resources and only result in a small number of hits [2]. Attention has been drawn to powerful generative deep learning techniques for efficient drug design due to their ability to learn a distribution for a given dataset and produce novel synthetic data from it. Deep generative models can explore chemical space efficiently through formulating a mapping between desired properties and corresponding molecular structures [3]. Attributed to the success of syntactic pattern recognition in language modelling tasks, early molecular generative models are designed on the basis of SMILES, a line notation that describes chemical structures using ASCII strings. However, SMILES representation is limited by its intrinsic instability in describing molecular structural information, as structurally similar molecules can result in dissimilar SMILES strings [4].

Graph-based generative models are considered particularly suitable for molecular generation, as molecular structures are naturally well-represented by graphs [5]. With success in applications of generative models and Graph Neural Networks (GNNs), many popular generative architectures such as Variational Autoencoder (VAE) [6], Flow-based models [7], are adapted to graph data structures and applied on molecule generation tasks. A common generation strategy starts with a latent representation with no atoms and bonds, building molecular graphs through autoregressive or one-shot generation. However, such generation strategy utilises random sampling to introduce novelty in the generated molecular structures. As molecule generation is extremely strict with precise validity and biological constraints, the basic graph-based molecular generation is unreliable and requires further control, which leads to the adoption of chemistry-aware constrained generation steps (e.g.[6]).

Fragment-based drug design (FBDD) approaches, which generate molecules conditioned on pre-existing sub-molecular units, constitute effective means to further constrain chemical space and improve generation of biologically active compounds [8]. Compared to random sampling schemes widely applied in exploring novel structures, using chemically meaningful fragments in molecular generation keeps search space chemically relevant and reduces the number of generation steps needed for frequent molecular motifs [9]. Moreover, FBDD strategies can utilise information obtained from structure-activity relationship analysis, and can preserve such useful biological activity properties in the synthesised molecules. FBDD strategies have been introduced and successfully applied in fragment-to-lead studies via different techniques, such as scaffold hopping [10], linker/PROTAC design [11], and R-group optimisation [7].

Optimisation approaches are often integrated with generative models to navigate through the search space and identify promising initial seeds which can generate molecules with specified properties. Such optimisation is usually implemented through heuristic optimisation techniques, such as, Bayesian Optimisation (BO) [12], or Reinforcement Learning (RL) [13]. However, these methods require additional computation while balancing exploration and exploitation within the explored space. For instance, BO may struggle with high-dimensional search spaces as the number of features increases and the search space becomes increasingly sparse, thereby requiring more time and training data to find promising regions to sample. RL is limited in multi-objective optimisation, as it requires large volume of data and involves a lot of computation. Whereas for molecular generation problem, the more conditions added to optimisation objective, the less suitable molecules are for policy training. Therefore, controlling multiple attributes in molecular design efficiently remains challenging.

To generate molecules that preserve chemically meaningful fragments and combine multiple attributes relating to different properties, we propose a controllable graph-based deep generative model for fragment linking, CLigOpt, towards target-specific ligand generation. CLigOpt takes two substructures with desired chemical functions and generates a linker that connect them, resulting in a valid molecule. CLigOpt incorporates a Controllable Generation Module (CGM), which evaluates an initial sample with respect to its chemical properties before generation, so that linkers satisfy multiple chemical property requirements. Our model exhibits consistent and strong performance across six evaluation metrics in both structural feasibility assessment and chemical usefulness. A case study on designing inhibitors for hDHFR is used to demonstrate applicability, and the quality of generated inhibitors is assessed via molecular docking simulations and retrosynthesis reaction estimation. The top prioritised molecules are provided as potential leads for hDHFR inhibition, together with the predicted retrosynthesis reactions.

The main contributions of our work are as follows: (i) a VAE-based model for linker design, where co-embeddings of nodes and edges are utilised to improve the information learned from molecular graphs, is developed and evaluated, (ii) a multi-objective controllable molecular generation module, CGM, is introduced to generate promising target-specific ligands, (iii) comparisons with state-of-the-art methods and multiple metrics are reported to evaluate results of the generative process.

## 2 Methodology

### 2.1 Data

The processed and filtered ZINC [14] and CASF [15] datasets were used as previously described [11] for general-purpose model training and testing (4174,997 examples from ZINC for training, 400 examples from ZINC and 309 examples from CASF for testing). Each example consisted of two fragments and a linker with size between 3 and 12 atoms. For target-specific linker design, 814 inhibitors and their binding affinities (pIC50) were obtained from ChEMBL [16], targeting human dihydrofolate reductase (hDHFR). The inhibitors were processed and fragmented following the procedure described in [17], resulting in 1576 fragment pairs, 80% of which were used for training affinity predictors, and 20% selected for target-specific controllable generation.

### 2.2 Model Framework

The proposed CLigOpt methodology is a graph-based Variational Autoencoder (VAE) framework which takes two molecular fragments as input and generates a linker to form a complete plausible molecule. CLigOpt contains two components, a generative network *θ* that maps from a latent representation *z* to a sample *x*, providing a joint distribution on

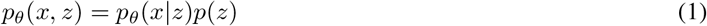

where *p*(*z*) is a smooth distribution, *p*_*θ*_(*x* | *z*) is the decoder that maps from *z* to *x*. An encoder *ϕ* maps from *x* to *z*, which is approximated by *q*_*ϕ*_(*z* | *x*) to reduce the computational cost. Our training and generative procedures are outlined in Figure 1 and Algorithms S1 and S2.

**Figure 1:**
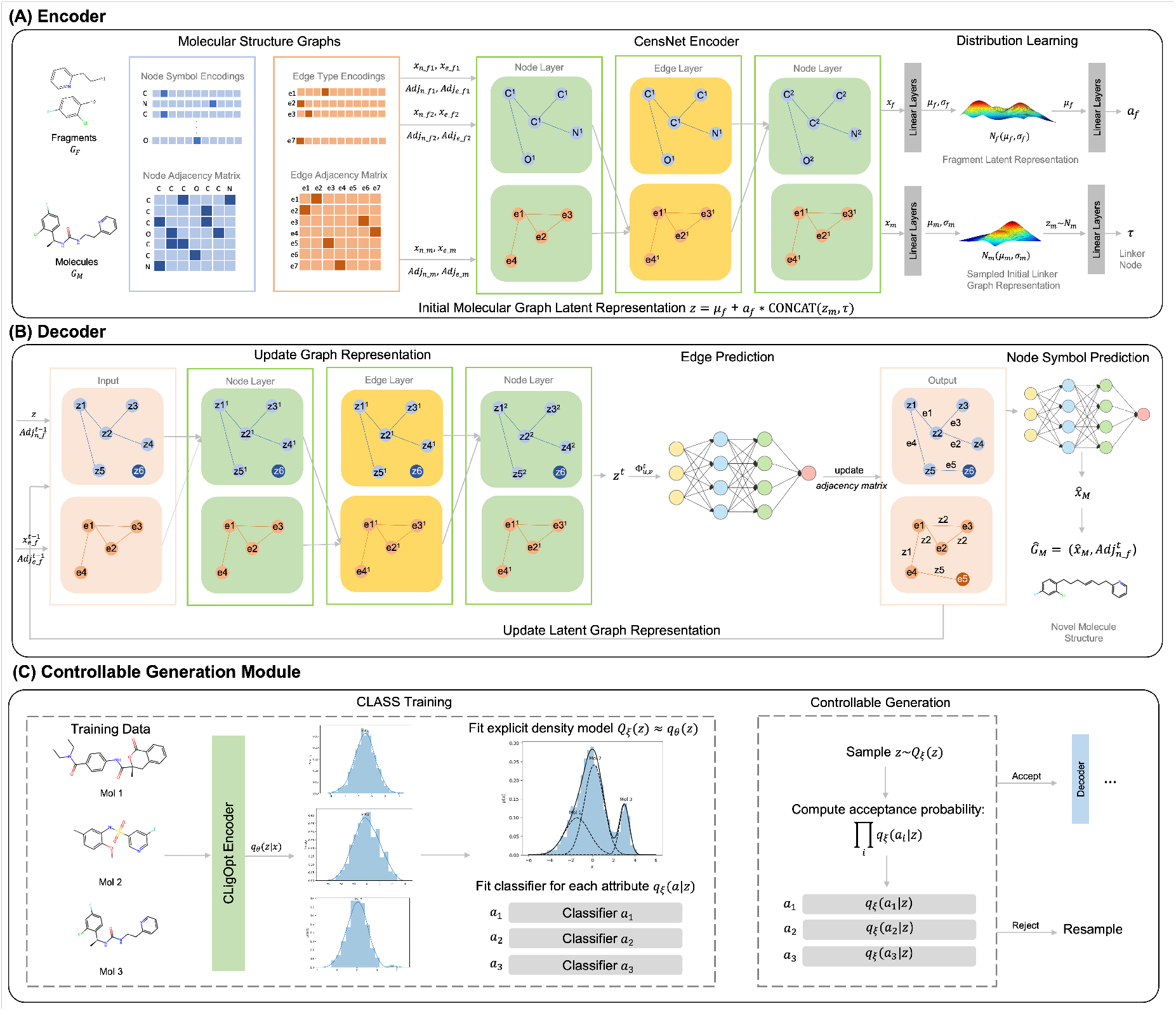
A diagrammatic representation of CLigOptCensNet. (A) shows an overview of the CensNet encoder, two fragments and a complete molecule are encoded into latent space, from where an initial representation for decoder is formed through the weighted sum of the average fragment representation and a sample is how the model Encoder encodes two molecules into the latent space, which is then decoded by our Decoder. (B) shows an overview of CensNet decoder, where the initial representation is applied through a sequential edge prediction process joining the two fragments. (C) the Controllable Generation Module (CGM) where our trained encoder is used to encode molecules into the latent space to fit an explicit density model and attribute classifiers, and then sample from the fitted density model and attribute classifiers before decoding each point into the final molecule.

#### Encoder

The workflow of encoder is shown in Figure 1 (A). Two models (GCN [18] and CensNet [19]) are employed as encoders and their ability to learn representations from hidden graph embeddings is investigated. Specifically, GCN is one of the most commonly used GNN architectures and its convolutions can be seen as message-passing through the node adjacency matrix, using only edge information to determine which nodes are adjacent to update node features. CensNet defines two types of layers, i.e. node and edge layer, which employ convolutions on line graphs, enabling the roles of nodes and edges to switch. Through convolution operations, co-embeddings of nodes and edges can be learned. Owing to this alternative updating rule, CensNet allows embedding nodes and edges to latent feature space simultaneously, as well as mutual enrichment over node and edge information.

We developed two version of CLigOpt models based on the two aforementioned networks. CLigOptGCN used three layers of GCNs as encoder, while the CLigOptCensNet encoder contained two node CensNet layers and one edge CensNet layer (see Figure 1 (A)). The means and variances, denoted as (*µ*_*f*_, *σ*_*f*_) and (*µ*_*m*_, *σ*_*m*_), of fragment *G*_*F*_ and molecule *G*_*M*_ are obtained by the encoder, which can be formulated as:

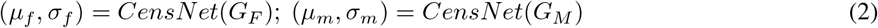

Then, the initial graph representation for decoder, *z*, is the sum of *µ*_*f*_ and the combination of *z*_*m*_ sampled from either the distribution *N*_*m*_(*µ*_*m*_, *σ*_*m*_) (for training) or *N* (0, 1) (for generation), and a latent node representation *τ* derived from *z*_*m*_ through a linear layer.

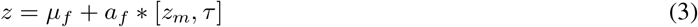

where [,] represents the operation of concatenation.

#### Decoder

Our training and generative procedures are explained in Algorithms S1 and S2. The decoder utilises a similar sequential generation scheme inspired by CGVAE [6] where GNNs are applied to predict edge connection through a bond-by-bond manner (see Figure 1 (B)). We utilise the same graph embedding network engaged in **Encoder** (GCN and CensNet) to learn an informative representation of the generated graph. Networks are improved by adding layer normalisation and dropout unit for each layer to reduce the impact of internal co-variate shift and prevent overfitting.

During generation, a set of nodes is initialised between the two fragments from which a linker is built through an iterative node update, edge selection and edge labelling process. At the beginning of each iteration, a focus node *u* is first selected for the decoder to predict its connectivity to the graph, including a pseudo stop node, subject to the valency constraints. The initial graph representation *z* is first be updated via a graph encoder layer. The graph representation *z*^*t*^ at the *t* − *th* step (iteration) is represented by,

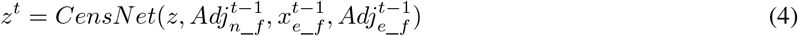

Then, for each remaining node *v*, a feature vector is constructed to predict the probability of an edge existing between *u* and *v* and the type of this edge. The feature vector integrates node-level information on the focus node and its edge candidates, as well as graph-level information for the initial graph representation and the current graph states. The feature vector for predicting the probability of an edge between *u* and *v* is provided by:

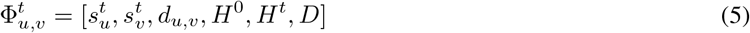

where 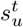 and 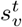 are the concatenation of hidden state *z*^*t*^ and atomic label representations of the focus node *u* and the remaining nodes *v* at step *t, d*_*v*,*u*_ is the graph distance between *v* and *u, H*^0^ is the average initial graph representation, *H*^*t*^ is the average representation of the current graph state, *D* represents the 3D structural information of the two input fragments, and the vectors inside square brackets are concatenated.

At the end of each step, the edge with the highest predicted probability is selected to connect to the focus node. The probability of an edge connecting to the stop node is also predicted through the same procedure. When the stop node is selected, the current focus node is finished and the nodes it connects then become the next nodes to focus. For step *t*, new status 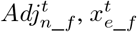 and 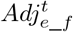 are updated by edge prediction module *MLP*_*E*_.

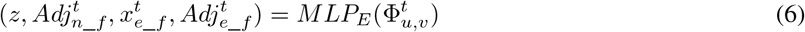

Such process is repeated until every remaining node is visited. The largest connected component is returned as the predicted molecule. After predicting all edges, the node symbols 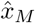 is generated by a node prediction module *MLP*_*N*_ .

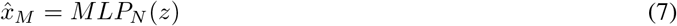

The reconstructed molecule *Ĝ*_*M*_ is then formed by 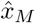 and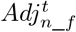 .

#### Training

Our model is trained with a standard VAE objective. For a given pair of fragments *G*_*F*_ and a molecule *G*_*M*_, the model is trained to reconstruct *G*_*M*_ from (*G*_*F*_, *z*) using Teacher Forcing [20], while enforcing the regularisation constraint on the encodings of *G*_*F*_ and *G*_*M*_ . Thus, the objective function comprises two components, a reconstruction loss and a regularisation loss:

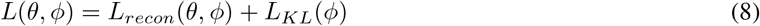

### 2.3 Target-Specific Controllable Generation

We built a Controllable Generation Module (CGM) referring to CLaSS [21] that optimises molecule generation for desirable properties. Given an initial sample *z*, and a set of independent attributes *a* ∈{0, 1} ^*n*^ = {*a*_1_, *a*_2_, …, *a*_*n*_}, CLaSS trains a set of classifiers that predicts the probability of *z* getting accepted on each attribute *a*_*i*_. The process of conditional generation of a molecule *x* given attributes the independent attributes, *p*(*x*|*a*), can then be formulated as:

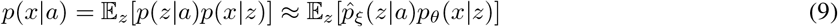

where 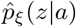 is derived using a density model *Q*_*ξ*_(*z*), in combination with a per-attribute classifier model *q*_*ξ*_(*a*_*i*_ | *z*), based on Bayes’ rule and conditional independence of the attributes, leveraging rejection sampling from parametric approximations to *p*(*z* | *a*). By enforcing multiple attribute constraints, the acceptance probability is the product of the scores from the attribute predictors, while sampling from the explicit density *Q*_*ξ*_(*z*).

#### Controllable Generation Module

We adapt CLaSS as our Controllable Generation Module, CGM, aiming to generate molecules with high drug-likeness, high synthetic accessibility and high binding affinity to a target protein. A schematic representation of our module is illustrated in Figure 1 (C). A Gaussian mixture density model is fitted over known molecules to estimate the posterior distribution of latent space *z*. From there, we design three classifiers in parallel, which take in a given latent embedding *z* and a pair of fragments encoded by CLigOpt, to evaluate the Quantitative Estimate of Druglikeness (QED), Synthetic Accessibility (SA) score, and the negative logarithm of IC50 (pIC50) respectively, aligning with the aforementioned three standards. Rejection sampling then operates through sampling from the fitted density model. CGM evaluates whether an initial sample is likely to satisfy certain standards on selected attributes through the previously mentioned classifiers, and thus decides whether to proceed with generation with this seed or not.

## 3 Results

The performance of CLigOpt was measured in different aspects of molecular generation, i.e. (i) fragment linking and forming plausible molecules, assessed through benchmark linker design (Benchmark Evaluation on Test Sets) and (ii) generating target-specific drug-like molecules (Controllable Target-Specific Ligand Generation), measured through a case study of generating lead candidates for hDHFR. Generated molecules were then prioritised and binding potential of the generated molecule set was evaluated via docking (Evaluation via Molecular Docking). Finally, synthesisability via specific reaction steps of promising molecules are indicated (Synthesisability of Generated Molecules).

### 3.1 Benchmark Evaluation on Test Sets

The ability of CLigOpt in generating reasonable molecules was evaluated on two benchmark datasets, namely ZINC and CASF, and compared with two baseline models for fragment linking, namely DeLinker [11] and FFLOM [7]. DeLinker is a VAE-based model that generates molecules in bond-by-bond manner, whereas FFLOM is a flow-based model that carries out one-shot generation of molecules. For each test set, 250 molecules were generated for each fragment pair, the quality of generated molecules was assessed on the basis of (i) structural usefulness via validity, novelty and uniqueness metrics, and (ii) chemical usefulness, i.e. Synthetic Accessibility (SA) score, ring aromaticity, and Pan-Assay Interference compounds (PAINS). The results are shown in Figure 2 and Table S1.

**Figure 2:**
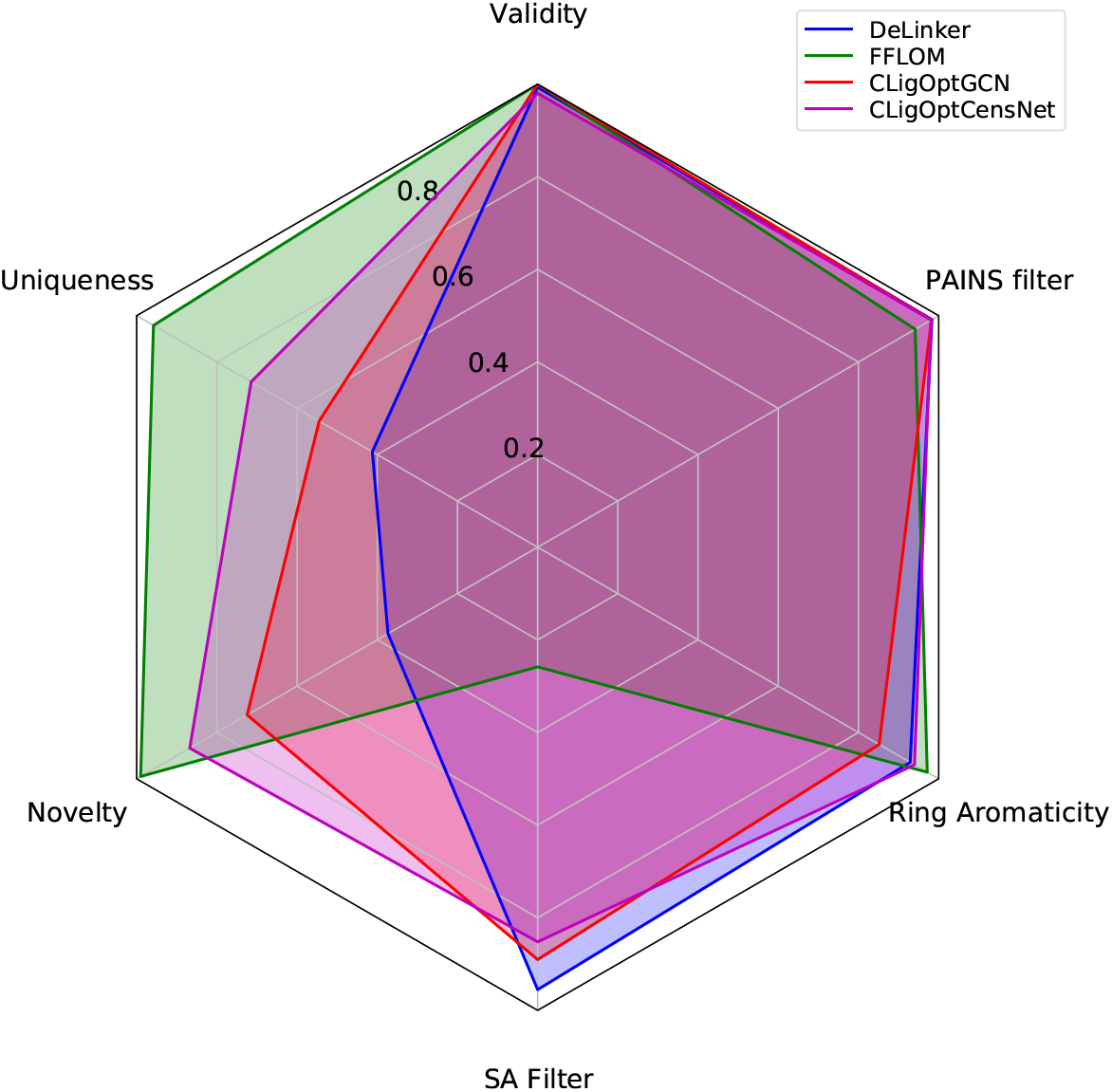
A radar plot showing the performance of our model as well as the two baselines. Although our model does not outperform the best model in each category, CLigOptCensNet achieves the best average performance.

#### Model Comparison

CLigOpt shows comparative but more consistent performance across all metrics compared to baseline models on ZINC test set. All models achieve similar performance in terms of validity, ring aromaticity and PAINS. CLigOptCensNet attains the best overall performance with high scores in every metric, while CLigOptGCN performs slightly worse in uniqueness. While DeLinker underperforms in uniqueness and novelty (30.21% and 49.44% less molecules compared to CLigOptCensNet), FFLOM underperfroms in SA filter (59.38% less molecules). Similar results are observed on CASF test set (see Figure S1), where CLigOpt outperforms DeLinker (58.42% on uniqueness, 48.58% on novelty) and FFLOM (49.34% on SA).

This indicates that DeLinker is able to generate molecules that are synthesisable and have good potential as drug candidates, but the generated set is repetitive and underperforms in novelty. On other hand, FFLOM can generate highly novel and unique molecules, however mostly difficult to synthesise. As a trade-off between these two cases, CLigOpt demonstrates competitive performance across all evaluation aspects.

### 3.2 Controllable Target-Specific Ligand Generation

A potential anticancer agent, hDHFR, is employed as target for a case study to investigate the ability of CGM to generate molecules with desired properties. QED, SA and pIC50 were used as the optimisation goals, aiming to generate molecular structures that fulfil pre-set conditions for three properties, (i) druglikeness, (ii) synthesisability and (iii) binding activity to hDHFR. The three conditions correspond to three classifiers predicting from an initial sampling of the latent space, to evaluate whether CLigOpt can generate a molecule with QED *>* 0.6, SA *<* 3, and pIC50 *>* 6. The classifiers can be used together or independently to generate molecules with different property control. We take the test set of hDHFR inhibitors described in Data, and generate 10 molecules for each pair of fragments for random sampling and rejection sampling.

#### Ground Truth Property Estimation

To evaluate the ability of CLigOpt in filtering out samples that did not meet the pre-set conditions, tools are employed to compute the true properties for the generated molecules as a guide. Due to concerns of introducing biases from overfitting, we utilise two pre-defined algorithms from rdkit [22] instead of neural networks, to compute the QED and SA score. Three machine learning approaches with diverse architectures were explored to predict pIC50: (i) transformer-CNN [23], a SMILES-based Quantitative Structure-Activity Relationship (QSAR) model that learns molecular embeddings from a transformer encoder, and predicts the corresponding activity using CharNN, (ii) modSAR [24, 25, 26], a two-stage QSAR model that involves detecting clusters of molecules, and applying mathematical optimisation-based piecewise linear regression to link molecular structures to their biological activities, and (iii) DeepAffinity [27], a semi-supervised deep learning model that utilises a unified RNN and CNN model to predict binding affinity. We compare Transformer-CNN and modSAR that are trained on a subset of hDHFR inhibitors, and DeepAffinity with the provided pre-trained parameters to select the most accurate model for predicting binding affinity.

Comparisons of the three models via the RMSE metric is shown in Table S5. Transformer-CNN outperforms Deep-Affinity and modSAR with RMSE = 0.82, and is selected to predict pIC50 for the generated molecules as a reference to evaluate the performance of CLaSS module. The results of the normalised fraction of molecules that meet the pre-set conditions, for a specific property as well as balancing all three properties, in a random generated set compared with CLaSS generated set are shown in Table 1. Evaluation on how CLaSS performs in a single property with different thresholds is shown in Table S2, S3, and S4.

**Table 1:** The proportion of molecules that pass certain filters when sampling randomly versus when sampling using the CLaSS accepted set. More results for each individual property can be seen in Tables S2, S3, and S4

|  | QED>0.6 | SA<3 | pIC50>6 | QED>0.6 &<br>SA<3 &<br>pIC50>6 |
| --- | --- | --- | --- | --- |
| GCN |  |  |  |  |
| Random | 9.78% | 16.27% | 53.32% | 1.48% |
| Accepted | 57.71% | 31.58% | 58.49% | 18.92% |
| CensNet |  |  |  |  |
| Random | 11.17% | 13.23% | 56.99% | 1.33% |
| Accepted | 49.47% | 22.07% | 62.78% | 10.35% |

#### Controllable Generation

Results show that CGM excels in efficiently generating more useful molecules. Table 1, S2, S3, and S4 report higher percentage of molecules meeting the pre-set conditions in CLaSS-generated set compared to the randomly sampled set, indicating that CLigOpt is able to identify the latent embeddings that result in molecules with desired properties. Table S2, S3, and S4 illustrates that, for QED and SA control, CLigOpt can generate a molecule set that contains three times more qualified molecules than random sampling. CLigOpt is less effective in pIC50 control, yet still better than random sampling. CLigOpt is also capable of multi-objective control, generating molecules that satisfy requirements for different goals. Moreover, the accepted set maintains a certain level of similarity to the original set in terms of QED and LogP (see Figure S2).

### 3.3 Evaluation via Molecular Docking

AutoDock docking [28] was employed, as developed by DeepChem [29], to further evaluate the binding of CLigOpt generated molecules. Molecular docking is conducted with the 3D structure of hDHFR on the CLaSS generated molecules with 10 highest-predicted pIC50. We compare the Binding Free Energy (BFE) of the generated molecules with 10 hDHFR inhibitors with highest binding affinities. Table 2 shows the results of average (E) and minimum (Min) BFE of these molecules, with respect to their binding affinities. Results indicate that CLigOpt is able to generate promising molecules that bind the druggable pocket of the 3D target structure with similar BFEs compared to known hDHFR inhibitors.

**Table 2:** Results summarising molecular dcking simulation.

| | Pred Aff | $E(kcal/mol)$ | Min |
| --- | --- | --- | --- |
| ChEMBL | 8.970 | $-8.606 \pm 0.711$ | -10.168 |
| GCN | 6.671 | $-8.256 \pm 0.581$ | -9.930 |
| CensNet | 7.058 | $-8.272 \pm 0.902$ | -9.732 |

### 3.4 Synthesisability of Generated Molecules

The number of reaction steps required to complete the synthesis of a given molecule provides an indication of its complexity with respect to commercially available materials. We randomly selected 100 molecules from CLigOptGCN and CLigOptCensNet generated set obtained in Controllable Target-Specific Ligand Generation, as well as the ChEMBL set as a baseline, to compute and compare the percentages of synthesisable molecules within each set. The retrosynthetic routes for each molecule were assessed using a Molecular Transformer-based [31] model [30], and it was observed that the generated molecules have higher successful retrosynthesis prediction than the ChEMBL set, with success rate of 84% molecules from the ChEMBL set, 96% from both generated sets. The distribution of the number of steps needed for each set is shown in Figure 3.

**Figure 3:**
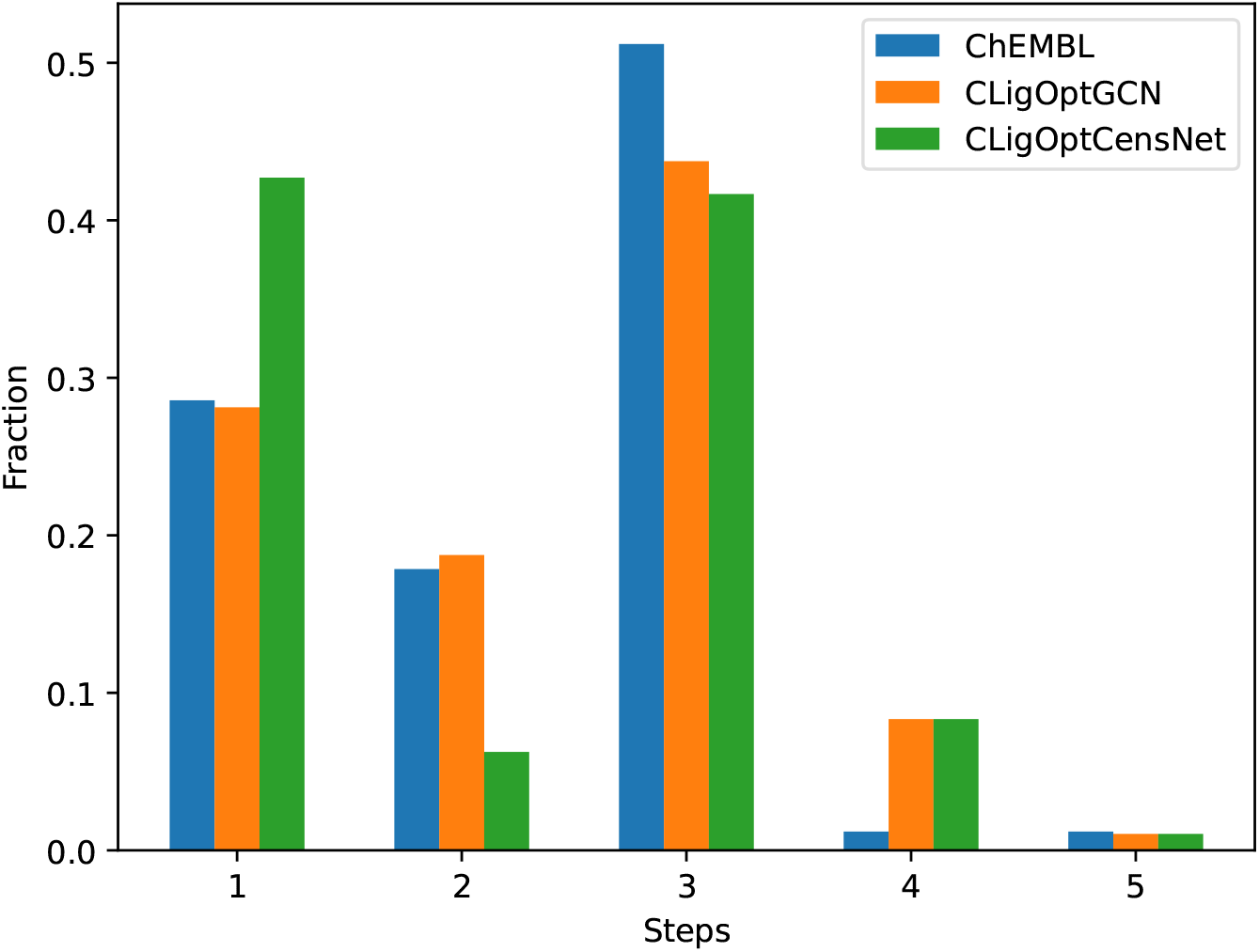
A histogram showing the proportion of molecules that could have a retrosynthetic path predicted by the model in [30]. Random samples of 100 molecules from each of our models and from 100 random molecules drawn from ChEMBL were used.

Overall, compared to the baseline set, CLigOpt generated molecules exhibit promising results in synthesisability regarding retrosynthetic analysis. Most molecules can be synthesised within 4 reaction steps. CLigOptCensNet-generated molecules show better synthesisability, managing most retrosynthetic routes in 1 or 3 steps. ChEMBL and CLigOptGCN generated molecules show similar results in the amount of molecules that can be made with in 2 steps, both achieving a peak at 3 steps. In comparison, more CLigOptGCN generated molecules require an additional step to synthesise than the ChEMBL molecules. We prioritise molecules from the two generated sets according to their QED, SA score, and pIC50, and provide the predicted retrosynthesis route for the best prioritised molecule in CLigOptCensNet in Figure 4.

**Figure 4:**
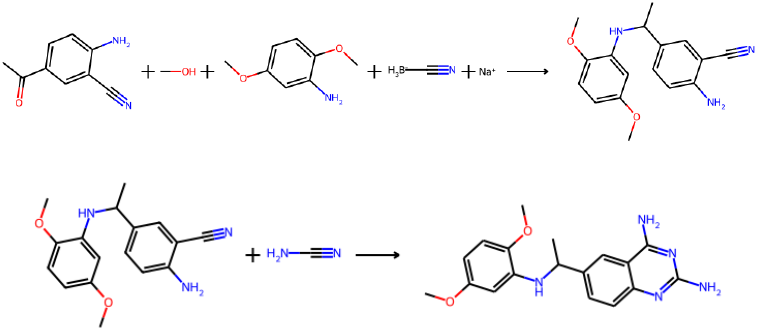
The two-step predicted retrosynthesis route for the best prioritised molecule (confidence: 0.92) in CLigOptCensNet. The molecule was prioritised according to its QED (0.655439), pIC50 (7.231879), and SA (2.643538)

## 4 Conclusion

In this work, a pipeline is introduced for efficient optimised molecule generation, CLigOpt, containing a Controllable Generation Module, where the generative process is restricted via rejection sampling. CLigOpt shows advantages over existing optimisation approaches in terms of reduced computational complexity, as it does not require heavy pre-training, surrogate model fitting, or policy learning. From comparing the quality of CLigOpt generated molecules with and without the property control module, we show that CLigOpt is an effective yet simple sampling scheme that improves the generation performance significantly and can handle multi-objective criteria well.

Evaluation of results is based on a variety of metrics that reflect the chemical and structural properties of the generated molecules. CLigOpt obtains consistent and strong performance across all six metrics. Performance of CLigOpt is evaluated with two encoders, GCN and CensNet. CLigOptCensNet shows slightly better performance than CLigOptGCN in novelty, uniqueness, and ring aromaticity, suggesting that employing co-embedding for nodes and edges provides more information than sole node embedding, and can also improve performance for the ensuing generative model.

The applicability of CLigOpt is illustrated via a case study on generating hDHFR ligands, evaluated through binding free energy simulation and retrosynthesis prediction. Generated molecules show similar binding compared to known hDHFR inhibitors, indicating the potential of CLigOpt to generate promising lead candidates. Future work can focus on a set of systematic optimisation objectives to further restrict and optimise searching molecular space.

## Acknowledgments

YL and XC are funded by the China Scholarship Council. PHdCA acknowledges funding through KCL/A*STAR (to MW and ST). MW is partially supported by A*STAR’s Decentralised Gap funding (I23D1AG081). ST acknowledges funding from the British Skin Foundation and the UK Royal Society.

## Notes

### Competing Interest Statement

The authors have declared no competing interest.

### Summary of Updates

Abstract, Introduction updated; Equations updated in Methodology; Figure 1 updated.

