## Supporting Information for "CLigopt: Controllable Ligand Design Through Target-Specific Optimisation"


**Yutong Li**
Department of Informatics
King's College London
UK

**Pedro Henrique da Costa Avelar**
Department of Informatics
King's College London
UK

**Xinyue Chen**
Department of Informatics
King's College London
UK

**Li Zhang**
Infocomm Research
A*STAR
Singapore

**Min Wu**
Infocomm Research
A*STAR
Singapore

**Sophia Tsoka**
Department of Informatics
King's College London
UK


**Algorithm S1** CLigOptCensNet: Training

**Input** Fragment graphs $G_F$, Molecular graphs $G_M$, and generation condition $c$
**Output** Edge probabilities $edge\_probs$, and $edge\_type\_probs$, and node probabilities $node\_probs$

$emb_F, emb_M \leftarrow Embedding(G_F), Embedding(G_M)$ ▷ Encoder
$H_F, H_M \leftarrow CensNet(emb_F), CensNet(emb_M)$
$H_{MG} \leftarrow GAP(H_M, batch)$
$\mu_F, \mu_M \leftarrow NN\mu(H_F), NN\mu(H_M), log(\sigma)_F, log(\sigma)_M \leftarrow NN_\sigma(H_F), NN_\sigma(H_M)$
$z_M \sim N(\mu_M, \sigma_M), noise \sim N(0,1)$ ▷ Adding fragment attention to the initial graph sample
$z_F \leftarrow Add(\mu_F, noise)$
$a_F \leftarrow Attention(z_F)$
$z \leftarrow z_F + a_F * z_M$
$z \leftarrow [z, H_M]$
**for** $t$ in $steps$ **do** ▷ Decoder
    $H^{(t)} \leftarrow CenNet(H^{(t-1)})$
    $edge\_probs^{(t)}, edge\_type\_probs^{(t)} \leftarrow NN_e(H^{(t)}), NN_{et}(H^{(t)})$
**end for**
$a_N \leftarrow Attention(z)$ ▷ Node Prediction
$H_N \leftarrow NN(z)$
$z_M \leftarrow H_N + a_N * z$
$node\_probs \leftarrow NN_{node}(z_M)$

**Algorithm S2** CLigOptCensNet: Generation

**Input** Fragment graphs $G_F$, and generation condition $c$
**Output** Result Molecule $mol$

$emb_F \leftarrow Embedding(G_F)$
$H_F \leftarrow CensNet(emb_F)$
$\mu_F \leftarrow NN\mu(H_F), log(\sigma)_F \leftarrow NN_\sigma(H_F)$
$z_M \sim GM(z), noise \sim N(0, 1)$
$z_F \leftarrow Add(\mu_F, noise)$
$a_F \leftarrow Attention(z_F)$
$z \leftarrow z_F + a_F * z_M$
$accept \leftarrow CLaSS(z)$
**if** not $accept$ **then**
    draw a new $z_M$
**end if**
$z \leftarrow [z, H_M]$
$edge\_probs, edge\_type\_probs, node\_probs \leftarrow NN_e(z), NN_{et}(z), NN_{node}(z)$
$mol \leftarrow$ assign $edge\_probs, edge\_type\_probs, node\_probs$

| | ZINC | | | | CASF | | | |
|---|---|---|---|---|---|---|---|---|
| Model | DeLinker | FFLOM | CLigOptGCN | CLigOptCensNet | DeLinker | FFLOM | CLigOptGCN | CLigOptCensNet |
| Validity | 99.33% | 100.00% | 100.00% | 98.02% | 98.00% | 100.00% | 100% | 99.38% |
| Uniqueness | 41.23% | 95.81% | 54.49% | 71.44% | 15.41% | 96.10% | 64.44% | 73.83% |
| Novelty | 37.32% | 99.00% | 72.44% | 86.76% | 41.03% | 99.21% | 77.46% | 89.59% |
| SA filter | 95.93% | 25.83% | 89.03% | 85.21% | 92.33% | 17.70% | 72.05% | 67.04% |
| Ring Aromaticity | 92.90% | 97.15% | 85.17% | 93.97% | 90.83% | 94.99% | 73.83% | 82.09% |
| PAINS filter | 98.32% | 94.14% | 98.20% | 98.12% | 99.41% | 92.85% | 97.33% | 97.92% |
| Average | 77.51% | 85.32% | 83.22% | **88.92%** | 72.84% | 83.48% | 80.85% | **84.98%** |

Table S1: Results for the two baselines (DeLinker and FFLOM) and our two models (CLigOpt{GCN,CensNet} for each of our evaluated metrics, as well as the average result along all models, on our two datasets (ZINC and CASF). We see that our CensNet model has the best average performance on both datasets.

| | QED>0.4 | QED>0.5 | QED>0.6 | QED>0.7 |
|---|---|---|---|---|
| CLigOptGCN | | | | |
| Random | 21.27% | 14.68% | 9.78% | 4.84% |
| Accepted | 73.75% | 63.99% | 57.71% | 28.91% |
| CLigOptCensNet | | | | |
| Random | 25.16% | 16.65% | 11.17% | 6.17% |
| Accepted | 55.26% | 52.00% | 49.47% | 28.09% |

Table S2: The proportion of sampled molecules that passed the QED filters for our models when sampling randomly versus when sampling using the CLaSS accepted set.

| | SA<3 | SA<2.5 |
|---|---|---|
| CLigOptGCN | | |
| Random | 16.27% | 2.66% |
| Accepted | 31.58% | 13.81% |
| CLigOptCensNet | | |
| Random | 13.23% | 3.99% |
| Accepted | 22.07% | 12.08% |

Table S3: The proportion of sampled molecules that passed the SA filters for our models when sampling randomly versus when sampling using the CLaSS accepted set.

| | pIC50>6 | pIC50>6.5 | pIC50>7 |
|---|---|---|---|
| CLigOptGCN | | | |
| Random | 53.32% | 24.27% | 11.49% |
| Accepted | 58.49% | 26.86% | 12.34% |
| CLigOptCensNet | | | |
| Random | 56.99% | 26.39% | 11.08% |
| Accepted | 62.78% | 29.02% | 12.17% |

Table S4: The proportion of sampled molecules that passed the pIC50 filters for our models when sampling randomly versus when sampling using the CLaSS accepted set.

| | Transformer-CNN | modSAR | DeepAffinity |
|---|---|---|---|
| RMSE | 0.8160 | 1.6902 | 1.7138 |

Table S5: Binding Affinity Predictor

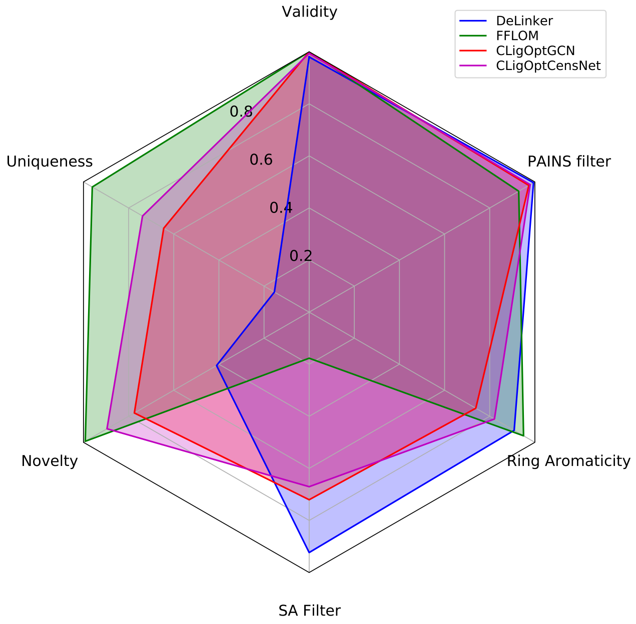


Figure S1: A radar plot showing the performance of our model as well as the two baselines on CASF. Although our model does not outperform the best model in each category, CLigOptCensNet achieves the best average performance.

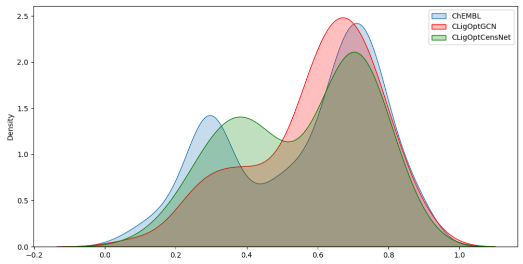


(a) The QED distribution of hDHFR inhibitors, molecule sets generated by CLigOptGCN, and CLigOptCensNet.

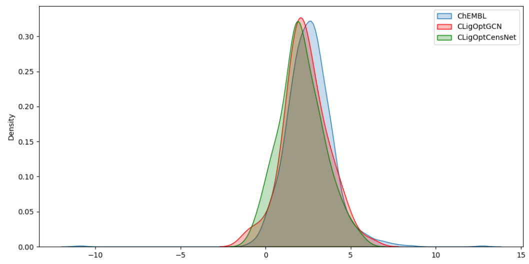


(b) The LogP distribution of hDHFR inhibitors, molecule sets generated by CLigOptGCN, and CLigOptCensNet.

Figure S2: The distribution of comparing attributes of generated sets and hDHFR inhibitors.
